# Early inflammation, immunopathology and aging

**DOI:** 10.1101/076828

**Authors:** Imroze Khan, Deepa Agashe, Jens Rolff

## Abstract

Age-related diseases are often attributed to immunopathology, which results in self-damage caused by an inappropriate inflammatory response. Immunopathology associated with early-life inflammation also appears to cause faster ageing, although we lack direct experimental evidence for this association. To understand the interactions between ageing, inflammation and immunopathology, we used the mealworm beetle *Tenebrio molitor* as a study organism. We hypothesized that phenoloxidase (PO), an important immune effector in insect defence, may impose substantial immunopathological costs by causing tissue damage, in turn accelerating aging. In support of this hypothesis, we found that RNAi knockdown of PO transcripts in young adults reduced inflammation-induced autoreactive tissue damage to Malpighian tubules, and increased adult lifespan. Our work thus provides empirical evidence for a causative link between immunopathological costs of early life inflammation and faster ageing. We also reasoned that if natural selection weakens with age, older individuals should display increased immunopathological costs associated with an immune response. Indeed, we found that while old infected individuals cleared infection faster than young individuals, they displayed exacerbated immunopathological costs and higher post-infection mortality. RNAi-mediated knockdown of PO response reduced immunopathology in older beetles and increased their lifespan after infection. This is the first demonstration of a direct role of immunopathological consequences of immune response during ageing in insects. Our work is also the first report that highlights the pervasive role of immunopathology under diverse contexts of aging and immune response.

## INTRODUCTION

Immunopathology refers to self-damage caused by an over-reactive immune system in response to infection, and contributes to the pathology of several human diseases (1–3). In insects, fast acting but nonspecific inflammatory responses are efficient against invading pathogens, but they also lead to immunopathology (4). Cytotoxin-producing effector systems like the phenoloxidase (PO) response pathway can also act against host cells (4–6). However, the role of immunopathology in natural populations is still unclear, with little information on whether or to what degree immunopathology explains variation in host immune response or impacts fitness. It is important to address these gaps, because immunopathology due to inflammatory responses can have long-term implications. For instance, it was proposed that reduced inflammatory exposure during childhood may have contributed significantly to increased lifespan in modern human industrialized societies (7).

Recently, the impact of an early life immune response on faster ageing was studied experimentally in the model insect *Tenebrio molitor* (8). The study predicted that accelerated ageing is caused by immunopathological damage to Malpighian tubules, fluid secreting epithelia that are functionally analogous to the human kidney (9, 10). However, we lack direct experimental evidence for this hypothesis. The phenoloxidase (PO) pathway, a fast acting immune effector in insects (11), is a likely candidate to cause the observed tissue damage via cytotoxic intermediates in *T. molitor* (12). Previous studies have already established a link between an over-active PO response and increased mortality in *Drosophila melanogaster*. For instance, flies that carried mutations in *serpin*-encoded spn27A failed to inhibit over-expression of the PO response, and died faster due to excessive melanisation (13). Also, mutations that inactivates a signaling serine protease (CG3066) in fruit flies– required for activation of pro-PO – increased lifespan after *Streptococcus pnuemoniae* infection (14). Based on these results, we propose that the immunopathological damage to Malpighian tubules caused by the PO cascade results in accelerated aging after early immune response.

In most animals, ageing is associated with a progressive decline in immune function leading to increased mortality and morbidity (15–17). Yet recent experiments seem to contradict this. For example, in *D. melanogaster* different immune genes show age-specific up-regulation-older fruit flies increased the expression of antimicrobial peptides after infection (18) and harbored lower pathogen load (19). Similar results were reported for the red flour beetle *Tribolium castaneum*, where ageing enhanced multiple components of immunity such as hemolymph antibacterial activity and PO response (20). However, despite this increased immune response with age, older beetles are more likely to die from infection. This suggests a mismatch between immune function and the ability to survive after infection, and questions whether the observed age-related increase in immunity is beneficial. Instead, relaxed natural selection in aged individuals (21, 22) may result in deregulation of the immune system, increasing immunopathological costs. We hypothesize that immunopathology associated with such a deregulated (over-reactive) immune response can directly contribute to greater post-infection mortality with age.

To begin to understand the complex interplay between ageing, immune response and immunopathology, we conducted experiments with mealworm beetle *T. molitor*. We first investigated whether PO-mediated immunopathology during early life inflammation resulted in faster aging. We then tested whether older individuals experience increased immunopathological risk and whether this is also mediated by the same mechanisms as in early life inflammation.

## METHODS

All experiments were conducted with *T. molitor* females. We collected experimental beetles from an outbred stock population maintained at 30 ± 2°C and supplied with an *ad libitum* diet of wheat bran and rat chow, supplemented with apple every 3 days. For all live infections, we used *Staphylococcus aureus* strain JLA513, a tetracycline resistant strain known to cause persistent infections in *T*. *molitor* (23). To produce inflammatory responses without pathogenesis, we used peptidoglycans (Sigma) derived from the *S. aureus* strain described above. We collected early pupae and determined their sex by observing the terminal abdominal segments. Upon eclosion, only females weighing between 120 and 150 mg were retained individually as virgins in grid box containers. All experimental females were thus controlled for age, mating status, and size. We used 7-day-old and 49-day-old individuals as ‘young’ and ‘old’ adults, respectively. We subjected both young and old females to experiments on the same day to enable a direct comparison between age classes.

### Experiment 1: Impact of immunopathology associated with early immune activation on ageing

#### (a) RNAi

To investigate whether reduced immunopathology minimizes the survival costs of early inflammation, we experimentally manipulated the degree of immunopathology during early life immune response and quantified survival. We used RNA interference (RNAi) to knock down expression of two proPO genes in 7-day-old virgin females: a previously sequenced *Tenebrio* proPO gene transcript (NCBI accession-AB020738.1; henceforth PO1 beetles) (24), and an ortholog to *Tribolium castaneum* proPO subunit 1 (NCBI accession-NP_001034493.1; henceforth, PO2 beetles). We amplified the internal region of the cDNA sequences encoding PO with T7-tailed primers (Table S1), and used them as templates to synthesize dsRNA using the T7 MEGAscript kit (Ambion) according to the manufacturer’s instructions. We extracted the resulting RNA with phenol/chloroform, re-suspended it in sterile nuclease free Ringer solution (128mM NaCl, 18mM CaCl_2_, 1.3mM KCl, 2.3mM NaHCO_3_ (23)), and stored it at - 80°C until further use. We annealed the complementary strands by heating at 65°C for 30 min and incubating at 22°C for an hour, before injecting 100ng purified dsRNA into each beetle. As RNAi control (or immune challenged control) (EI), we injected individuals with an internal region of the cDNA sequence encoding a lysozyme (Gm-Lys) from *Galleria mellonella* (Swiss-Prot accession-P82174), available in our lab (25). We monitored RNAi efficacy by performing qPCR (see supplementary information for qPCR methods and primers; Table S2). We used the comparative CT method (26) to estimate relative gene expression (see Fig. S1 & Table S3 for RNAi efficacy).

Two days after RNAi, we injected each beetle with 5µl peptidoglycan (concentration: 100ng in 1 ml of Ringer solution) to deplete the basal amount of PO in the hemolymph. Finally, after another 2 days, we challenged each beetle with a higher dose of peptidoglycan (5 µl; 5µg in 1 ml of Ringer solution) to induce a strong immune response. Following the immune challenge, a subset of beetles from all RNAi treatments (n = 30 beetles/ treatment) were monitored for total lifespan. Four days after immune challenge, the remaining beetles were either assayed for Malpighian tubule (MT) activity (n = 20-28 beetles/treatment) or phenoloxidase response (n = 16 beetles/treatment) (see below for methods). A set of 30 beetles served as unhandled full control (Control) that remained in the grid box container throughout the experimental window. In addition, as a procedural control (PC) for the impact of early-immune activation on adult life span, 30 beetles received ds-Lys injection (mock RNAi) followed by a mock immune challenge (with insect Ringer). Below, we briefly describe the methods for quantifying immunopathology and PO response.

#### (b) Malpighian tubule activity

A prior study by Sadd and Siva-Jothy (12) used a modified ‘oil drop’ technique (27, 28) to demonstrate a large reduction in Malpighian tubule (MT) function due to immunopathology associated with immune induction. The method provides an *in vitro* functional estimate of the ability of isolated tubules to transport saline across the active cell wall into the tubule lumen. We estimated the fluid transporting capacity of MTs harvested from experimental beetles 4 days after immune challenge, as a proxy for immunopathology due to immune response. Each beetle has three pairs (dorsal, lateral and ventral) of large MTs of varying length but similar secretion rates (29). Hence, we dissected one tubule from each cold-anaesthetized animal under cold sterile modified *Tenebrio* Ringer saline, prepared as described in Wiehart et al. (30). We severed the tubule at the point where it connects to the gut, and removed another ~0.5mm length from the open end (to control for the condition of the cut end). Following this, we transferred a single tubule per beetle to a 60 µl drop of sterile modified Ringer saline supplemented with 0.05% w/v phenol red to facilitate visualization, and 0.1mM L^-1^ dibutyryl cyclic AMP to stimulate fluid secretion (12, 28). We covered the whole preparation with mineral oil (Sigma). Next, we pulled the open end of the tubule out of the saline drop and wrapped it around 0.1 mm pins (Fine Science Tools) in the mineral oil, where it secreted fluid. 6 hours later, we measured the volume of the secreted droplet, as well as the length of tubule that remained within the saline drop using ImageJ software. The volume of the secreted droplet is negatively correlated with the degree of immunopathological harm to Malpighian tubules.

#### (c) Phenoloxidase response

We measured the PO activity of RNAi-treated beetles by measuring the rate of formation of dopachrome with a spectrophotometer (31). We mixed 2 μl undiluted hemolymph (collected from a wound between the head and thorax) with 8 μl PBS, and centrifuged the sample at 6500 rpm for 15 minutes at 4°C. We transferred 5 μl of the supernatant to a 96-well-microplate containing 20 μl PBS and 140 μl distilled water to measure activated PO enzyme (henceforth, PO activity). We then added 20 μl of L-Dopa substrate into each well, and transferred the plate immediately into a Microplate reader. We allowed the reaction to proceed at 30°C for 40 minutes, and then measured absorption at 490 nm once every minute. We quantified PO enzyme activity as the slope of the linear phase (between 15 to 45 minutes) of the reaction in each well (Vmax: change in absorbance per minute).

### Experiment 2: Impact of ageing on immune function and post-infection survival

We grew a *S. aureus* culture overnight in liquid LB medium to an OD_600_ of 95%. We then centrifuged the culture, washed the pellet three times before re-suspending in insect Ringer solution. We injected 5 μl of this suspension directly into the haemocoel of each individual (approximately 4 × 10^6^ CFUs-colony forming units-per inoculum; see (23)). Control individuals were injected with 5 μl of Ringer solution. After infection, we redistributed beetles individually into grid boxes under standard conditions with access to food. For a subset of beetles (n = 15-17 beetles/ age group/treatment), we monitored individual survival daily at 8pm for 40 days. Another subset was tested for clearance of *S. aureus* infection from hemolymph after 6 hours, 1 day, 7 days and 14 days. At each time point, we harvested bacterial cells from a group of 9-11 individuals to estimate remaining CFUs per beetle using the perfusion bleeding method described by Haine and colleagues (23). Briefly, perfused hemolymph was collected from each beetle and plated on LB agar containing 5µg/ ml tetracycline as a selective agent. The number of colonies observed after 48 hours of incubation at 30°C should be negatively correlated with the ability to clear bacterial infection.

Next, we tested whether induced hemolymph antimicrobial activity against *S. aureus* cells differed across beetle age groups (n = 9-13 beetles/ infection treatment/ age group), using an *in vitro* cell killing reaction as described in Haine et al. (23). We collected 2 μl of undiluted hemolymph sample as described above, diluted it with 48 μl PBS and 2 μl of an overnight culture of *S. aureus* (approximately 10^6^ CFU), and incubated at 30°C with shaking at 150 rpm for 2 hours. Following this, we diluted the mixture 800 times and plated out as described above. The number of CFUs that appeared was counted after 48 hours incubation at 30°C. The number of colonies observed was used as a measure of (inversely related to) the strength of induced anti-S *aureus* activity of the beetle hemolymph.

Finally, we measured PO response (n = 30 beetles/age group) and estimated relative expression of the antimicrobial peptide-coding genes *attacin 2* and *tenecin 1* (n = 5-6 pairs/ age group/ gene; see Table S2 for qPCR primer sequences) as a function of age in naïve beetles as described earlier. Since these assays were performed with uninfected beetles, they served as an estimate of age-associated changes in baseline constitutive innate immune function in the absence of immune induction.

### Experiment 3: Impact of ageing and infection on immunopathology

Beetles from both age groups (7-day-old vs. 49-day-old) were first infected (or sham-infected) as described earlier (n = 11-14 beetles/ age-group/ infection status). Four days later, we estimated MT function as a proxy for immunopathology (see experiment 1 for methods). We also manipulated the impact of bacterial infection on MT activity and tested whether reducing damage to MTs can rescue the low post-infection survival of older beetles. To this end, we injected 45-day-old beetles with 5 µl of dsRNA (100ng/µl) of PO1 or Lys (n = 27-34 beetles/ RNAi treatment) as described in experiment 1. Two days later, we infected beetles as described above. We monitored a subset of individuals for their post-infection survival daily at 8 pm for 25 days (n = 15-18 beetles/ RNAi treatment). The remaining individuals were tested for MT activity as described above to quantify immunopathology (n = 12-16 beetles/ RNAi treatment).

## Data analysis

Residuals of bacterial clearance data were not normally distributed (tested with Shapiro-Wilks test). Hence, we log-transformed the data, and confirmed that the transformed residuals were normally distributed. Following this, we used a two-way or one-way ANOVA to test the following effects: (a) Bacterial clearance as a function of age and assay time (b) PO response of early-infected beetles as a function of RNAi treatments. We tested for pairwise differences between treatments after correcting for multiple comparisons, using Tukey’s HSD. Non-normally distributed data that could not be transformed to a normal distribution were analyzed using nonparametric Wilcoxon Rank Sum tests: (a) MT activity after early-immune response as a function of RNAi treatments (b) antimicrobial activity as a function of age (analyzed separately for sham infected and infected beetles) (c) PO response as a function of age (d) MT activity as a function of age and infection (e) MT activity of older beetles as a function of RNAi treatments. Here, we used a Steel-Dwass test to estimate pairwise differences.

We used Cox Proportional Hazard survival analysis to test the following effects: (A) beetle survival after infection as a function of age and infection (B) post-infection survival of old beetles as a function of RNAi treatments. We did not have any censored values while analyzing the data, as all beetles died within the experimental window. We calculated the impact of treatment (e.g. infection or RNAi) as the estimated hazard ratio of experimental vs. control group (hazard ratio = rate of deaths occurring in experimental group/ rate of deaths occurring in control group). A hazard ratio significantly greater than one indicates an increased risk of mortality in the experimental group compared to control individuals.

We analyzed median and maximum lifespan to measure the change in ageing rate following early-life immune activation. We used accelerated failure time (AFT) models (32) to examine the median lifespan in R using the ‘survival’ package, with each model separately analyzing the difference in median lifespan between a treatment and unhandled control group. We also compared knockout (PO1and PO2) and early immune challenged control (EI) beetles to analyze whether PO knockdown increased lifespan after an early immune response. We found that the Weibull distribution minimized the AIC value (Akaike’s Information Criterion) of AFT models, and was thus most appropriate to use for each comparison. For each model, we estimated the c-parameter (exponential (estimated coefficient associated with lifespan)) representing the difference in median lifespan between the two groups, as suggested by Swindell (32). A c-parameter value significantly less than 1 indicates reduction in lifespan and vice-versa. For a given comparison, the value 100(c −1) represents the percent change in median lifespan of the experimental vs. control group (33).

Maximum lifespan has been suggested as an important indicator of the ageing process (34); hence, we used it to test the impact of early immune response on ageing. We first estimated the 90th percentile lifespan when all treatments were combined, and then calculated the percentage of beetles in each treatment group living until this time. We then performed exact unconditional tests using a contingency table approach to compare percentage survival of each treatment group with unhandled control beetles (www.stat.ncsu.edu/exact). We also compared RNAi-treated beetles and immune challenged control beetles (i.e. PO1 or PO2 vs. EI) to test whether PO knockdown increased maximum lifespan significantly. We used a binomial, two-way model to generate a pooled z-score test p-value for each comparison. To obtain the treatment effect for each comparison, we divided the percentage survival of experimental groups by that of the respective controls.

## RESULTS

### Phenoloxidase response after early inflammation increases mortality and organ damage

We found that an early immune challenge in young adults caused an increase in PO response (Fig. 1A, Table S4A), with a concomitant reduction in MT activity (Fig. 1B, Table S4B). We further observed that RNAi knockdown of the PO response reduced MT damage in immune-challenged beetles, resulting in a limited reduction of fluid secretion rate (compare Fig. 1A & B, Table S4A & B). An AFT model showed that early immune challenged beetles also had shorter lifespans compared to full control and procedure control beetles (Figures 2A & B, Table S5A). We did not detect mortality until 16 days following the immune challenge, suggesting that early immune challenge did not have an immediate impact on survival (Figure 2A). The c-parameter was lowest in immune challenged control (EI) beetles, suggesting a reduction in median lifespan (percent decline with respect to full control, 100(c-1): ~35%) following an early-life immune response (Figure 2B, Table S5A). The negative impact of an early immune challenge on beetle lifespan could also be reversed by RNA interference of proPO transcripts (e.g. PO1 and PO2) in immune challenged beetles (Figure 2A & B, Table S5A). RNAi of both the transcripts extended the life span by ~31-32 % compared to immune challenged control (EI) beetles (Figure 2B, Table S5A).

**Figure 1.**
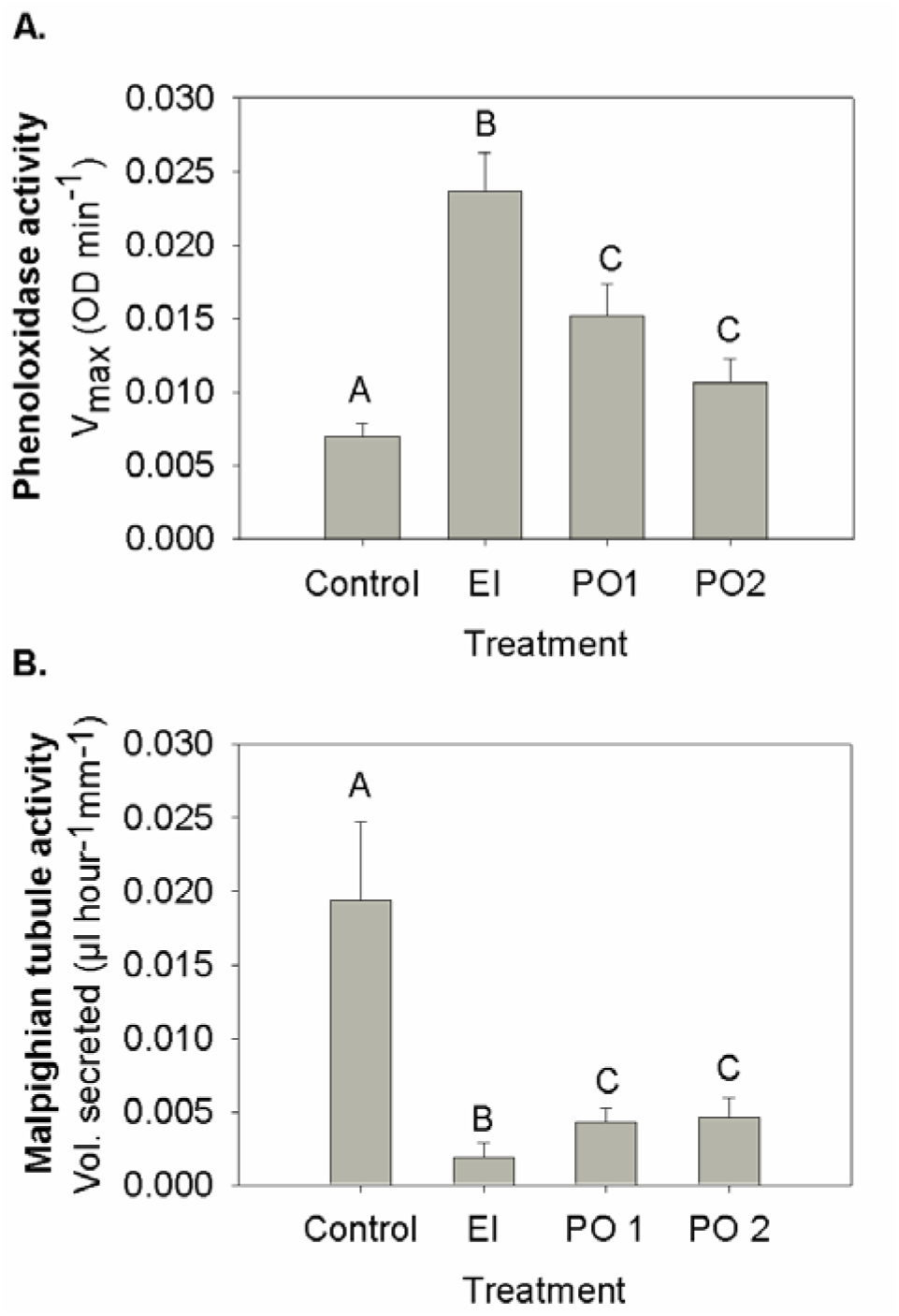
Impact of RNAi-mediated knockdown of pro-phenoloxidase transcripts on (A) phenoloxidase (PO) response and (B) Malpighian tubule activity, serving as a proxy for immunopathological damage, measured at day 4 post-immune challenge (i.e. day 11 post-emergence). PO activity was measured as Vmax of the enzymatic reaction with L-DOPA substrate. Malpighian tubule activity was measured as the rate of fluid secretion. Significantly different groups are indicated by distinct alphabets (based on Tukey’s HSD/ Steel-Dwass test). Control= Unhandled control beetles, EI= Early immune challenged control beetles, PO1= RNAi of PO1 transcript followed by an early-inflammation, PO2= RNAi of PO2 transcript followed by an early inflammation.

**Figure 2.**
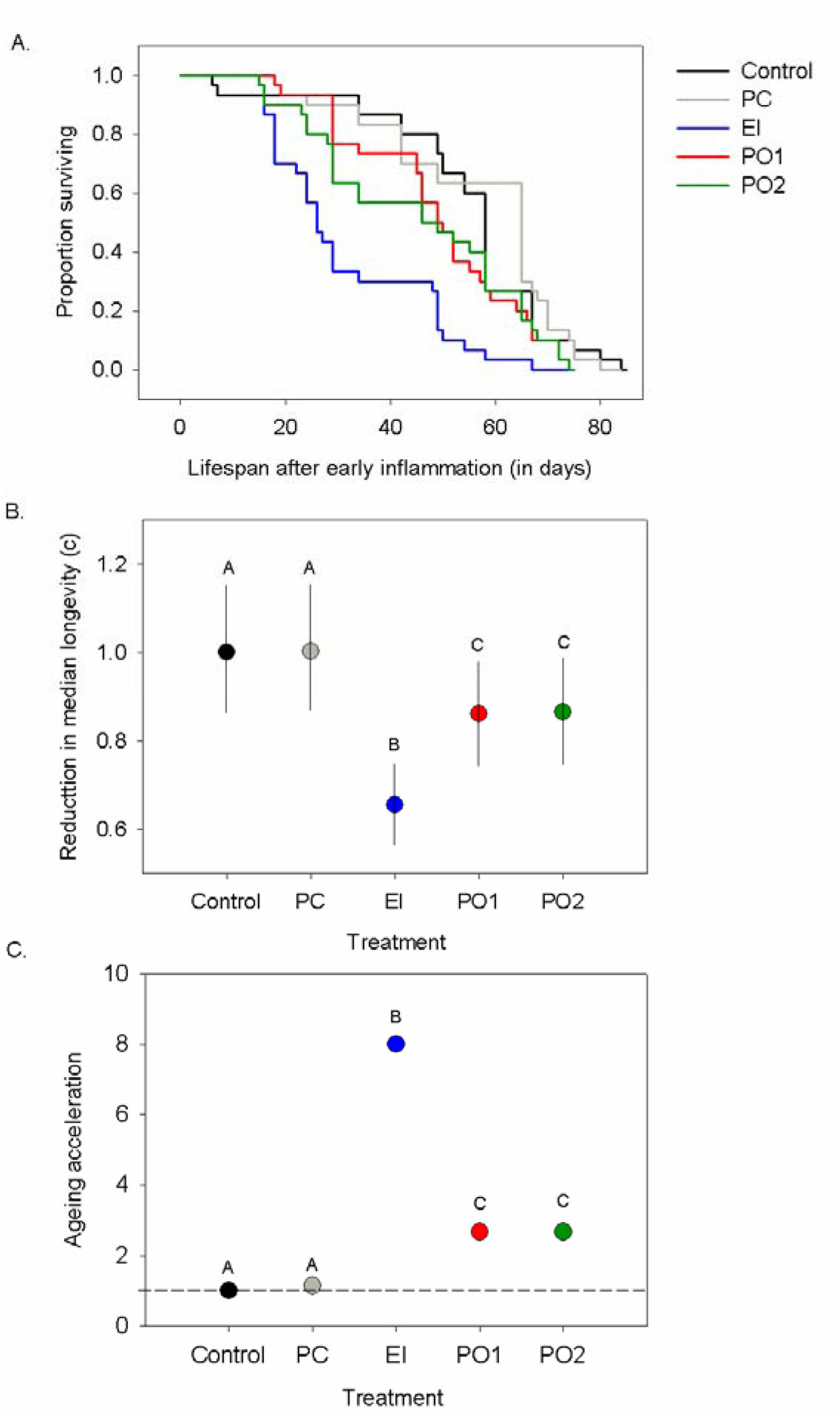
**(A)** Survival curves of adults after an early immune challenge that induced inflammation. **(B)** Impact of phenoloxidase knockdown on median lifespan. Survival of each experimental group was compared with the unhandled control group using an accelerated failure time model. The c-parameter denotes the effect of the immune challenge treatment on survival, averaged over total survival time. Error bars represent 95% confidence intervals. **(C)** Impact of phenoloxidase knockdown on maximum lifespan. ‘Ageing acceleration’ on the y-axis denotes the reduction in maximum lifespan caused by immune challenge treatments (the percentage of survivors to the 90th percentile survival time in the full control group/ the percentage of survivors to the 90th percentile survival time in the experimental group). Control = Unhandled control beetles, PC= Procedural control, EI= Early inflammation control beetles, PO1= RNAi of PO1 transcript followed by an early inflammation, PO2= RNAi of PO2 transcript followed by an early inflammation.

Analysis of maximum lifespan produced similar results as that of median lifespan. The 90th percentile of overall survival time was 74 days (pooling all individuals across treatments); the percentage of immune challenged individuals surviving to this time was significantly lower than control groups (EI = 3%, PO1 = 10%, PO2 = 10%, PC = 23%, Control = 27%; also see Figure 2C & Table S5B for treatment effects). These data suggest acceleration in ageing due to early-life immune response (compare 90^th^ percentile survival for each experimental group: FC = 79.5; PC = 83.1, EI = 57.6; PO1 = 73.8, PO2 = 73.8). Finally, we found that RNAi knockdown of the PO response significantly increased maximum lifespan compared to immune challenged EI beetles with normal PO levels, suggesting delayed ageing in knockout groups (compare ageing acceleration in Figure 2C, Table S5B).

### Immune responses increase with age

We found a rapid clearance of bacterial cells in both young and old beetles: *~*98% cells were removed within 6 hours after infection. However, older individuals showed consistently lower bacterial loads across different time points (i.e. 6 hours, 24 hours, 7 days and 14 days) after infection (Fig 3A, Table S6A). Next, we tested the ability of cell-free hemolymph to kill *S. aureus* cells 1 or 7 days after infection with live bacteria (or sham infection). We found that hemolymph from older infected beetles showed a significantly higher antibacterial response compared to younger beetles (Fig 3B, Table S6B). In contrast, age did not have a strong influence on the antibacterial activity of cell-free hemolymph from sham-infected beetles (Figure S2, Table S6B). Older beetles also showed an enhanced PO response (Fig 3C, Table S6C) and higher expression of the antimicrobial peptides *attacin* and *tenecin 1* (Fig. 3D, Table S3), even in the absence of a previous bacterial infection.

**Figure 3.**
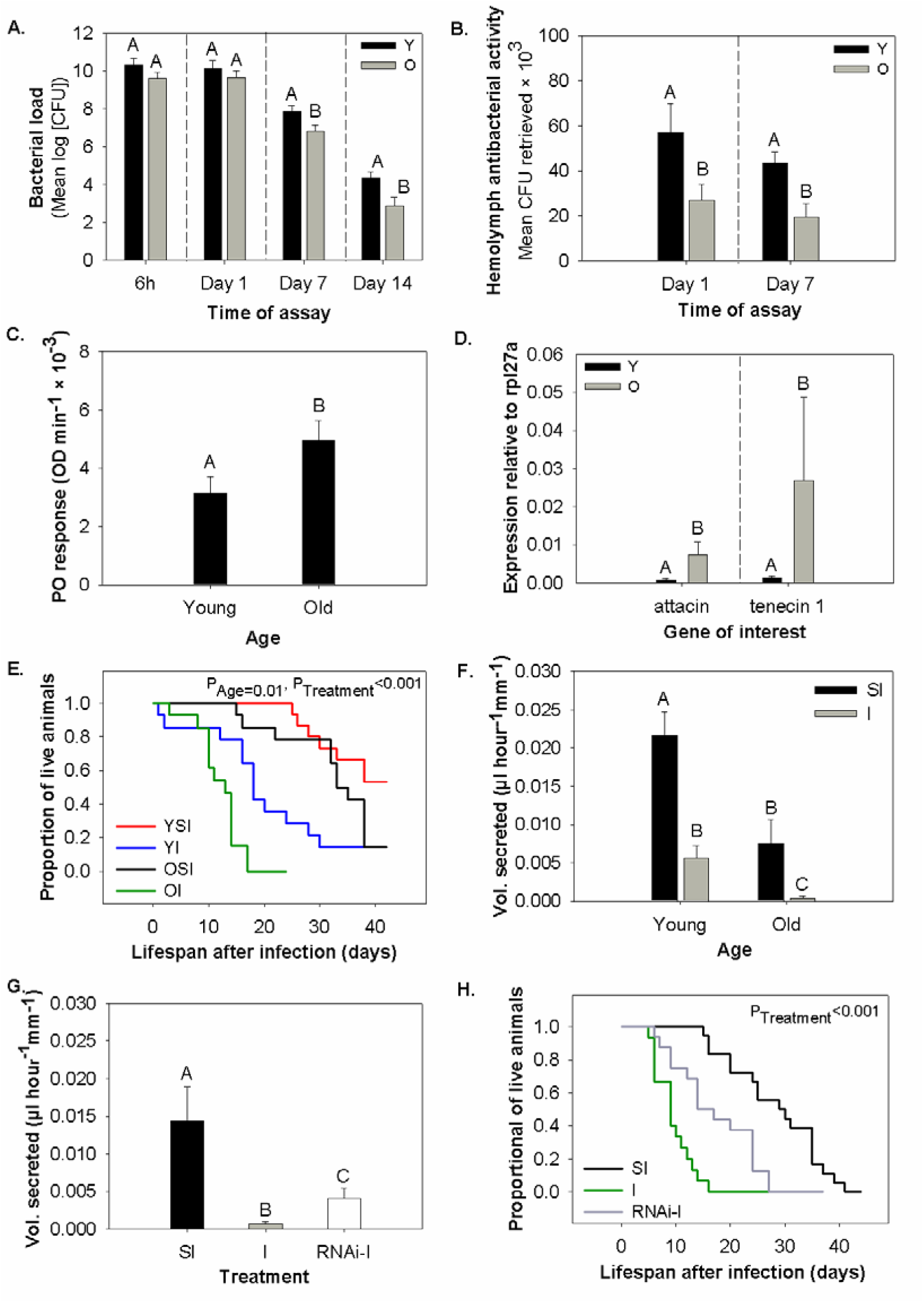
**(A-F)** Impact of age on (A) bacterial clearance at different time points after infection (B) hemolymph antibacterial activity of infected beetles at different time points (C) phenoloxidase (PO) response of naïve beetles (D) expression of antimicrobial peptide genes *attacin* and *tenecin-1* in naïve beetles relative to an internal control (*rpl27a*) (E) post-infection survival, & (F) Malpighian tubule activity after bacterial infection. **(G-H)** Effect of RNA interference of PO1 transcript on (G) immunopathological damage to Malpighian tubules, & (H) post-infection survival in older beetles. Bacterial clearance was measured as log_10_ of CFUs (colony forming units) recovered from perfused hemolymph after injection of *S. aureus* cells into each beetle. Hemolymph antibacterial activity was measured as the number of *S. aureus* cells recovered after 2 hours of exposure to beetle hemolymph. PO response and immunopathology were measured as described in Figure 1. Significantly different groups are indicated by distinct alphabets (based on Tukey’s HSD/ Steel-Dwass test). For panels A, B and D, alphabet assignments are meaningful only within each time point (or gene) (partitioned by dashed vertical lines), and are not comparable across time points (or genes). Y = Young beetles; O = Old beetles; YSI = Young sham-infected beetles, OSI = Old sham-infected beetles, YI = Young infected beetles, OI = Old infected beetles, PC = Sham infection with insect Ringer; I = *S. aureus* infection; RNAi + I = RNAi followed by *S. aureus* infection.

### Older beetles die faster after bacterial infection

To test if an age-related increase in immunity also confers greater survival benefits, we tested the impact of *S. aureus* infection on the survival of young vs. old beetles. Compared to young beetles, we found that old beetles died much faster after an infection (Hazard ratio: Sham-infection vs. infection in old beetles = 12.59, P<0.001; Sham-infection/ infection in young beetles = 5.18, P<0.001; Figure 3E, Table S6D). In contrast, beetle age did not alter the survival of sham-infected beetles significantly (Figure 3E, Table S6D).

### Increased immune response impairs Malpighian tubule activity via immunopathology

We found that ageing itself reduced the baseline MT activity in sham-infected beetles. Bacterial infection impaired the fluid transport ability of MTs in both age groups, but the relative impact of infection was much larger in infected older beetles (~95.1% reduction, compared to a 74% reduction in young beetles) (Fig. 3F, Table S7A). As observed for early immune-challenged beetles, we found that RNAi mediated knockdown of pro-PO1 partially rescued the MT activity of infected older beetles (Fig. 3G, Table S7B), and proPO1 knockdown beetles survived longer after infection than wild-type beetles (Hazard ratio: knockout vs. infected wildtype beetles = 0.396; P = 0.01) (Fig. 3H, Table S7C). We thus suggest that bacterial infection induced immune upregulation in older beetles, but incurred greater immunopathological damage to MTs, resulting in greater mortality despite higher immune function.

## DISCUSSION

In the work presented here, we evaluated the importance of immunopathology in the diverse contexts of ageing and immune response in the model insect *T. molitor*. We first show that early-life inflammation leads to faster ageing. Early-life immune activation (without the direct cost of a live and replicating pathogen) was sufficient to inflict immunopathological damage on Malpighian tubules (MTs), a vital insect organ that is functionally equivalent to the vertebrate kidney. Subsequently, we demonstrated that experimental suppression of the phenoloxidase (PO) response reduced the immunopathological damage to MTs and improved survival. These results highlight that a greater collateral damage to vital organs by PO is indeed one of the mechanisms that cause accelerated death of older beetles after early life inflammation. Although our work focuses on an insect system, it provides the first empirical support for the proposed link between childhood inflammation, immunopathology and reduced human lifespan (7). Next, we describe the immunopathological risk associated with an ageing immune system and its impact on the lifespan of older individuals. While ageing increased several aspects of beetle immunity such as PO response, antimicrobial peptide gene expression, induction of hemolymph antibacterial activity and the ability to clear *S. aureus* infection, it also severely impaired MTs and compromised the ability to survive after bacterial infection. We found that RNAi knockdown of the PO response once again minimised immunopathological damage to MTs of old beetles, partially rescued fluid transport ability and extended adult lifespan after infection. Thus, one of the most interesting implications of our work is the pervasive role of immunopathological damage to MTs that underlies the effects of early immune activation as well as the later impacts of infection.

In a prior study, Ayres and Schneider had predicted a direct role for PO response pathway in immunopathological damage and increased post-infection mortality of fruit flies (14). Our data not only support this prediction, but also reveal impaired MT activity as an important mechanism underlying the immunopathological consequences of the PO response and associated reduction in lifespan. We note that other components of the immune system can also be involved in immunopathology, and the PO pathway may not be the sole determinant of immunopathology. For instance, the Tumor Necrosis Factor-like protein encoded by the gene *eiger* is known to cause immunopathology in fruit flies (35). NF-kB signaling, which is involved in chronic inflammation in vertebrates (36), may also cause immunopathology in insects, though it is poorly studied. We thus suggest the need for further studies to test the fitness impacts of immunopathology caused by other immune effectors.

Another important finding of our study is a mechanism to explain the discordance between immune response and the host fitness post-infection (i.e. the ability to survive after bacterial infection), reported in this study as well as in flour beetles (20) and fruit flies (37, 38). Such a mismatch suggests that the inherent ability to mount an immune response is not a reliable predictor of host fitness after infection. Our data suggest that these contradictory results may be mediated via a large reduction in MT activity in infected older individuals. As a result, old beetles may pay greater immunopathological costs of increased immune activation compared to young beetles. In fact, several studies in vertebrates show that ageing leads to chronic inflammatory responses via maladaptive impacts of the innate immune system, contributing to the pathology of several age-related illnesses (17, 39). An exaggerated inflammatory response with age can induce lethal immunopathology in mice: old individuals die faster due to hepatocyte necrosis caused by an elevated level of interleukin-17 and neutrophil activation (36). Another study found increased post-infection mortality in older mice with suppressed anti-inflammatory cytokine interluekin-10 expression, suggesting a link between overactive inflammatory responses and increased risk of mortality (40). Thus, we suggest that the observed increase in beetle immunity with age represents immune activation at an unnecessarily high level (characteristic of pro-inflammatory profile with age in vertebrates) without any adaptive value. Instead, ageing leads to a dramatic increase in the negative impacts of inflammation, and the net impact of the immune response compromises old individuals’ ability to survive after infection. Ageing may also reduce tolerance to infection, the ability to limit negative impacts of pathogens, or self-damage (39). Consequently, older individuals may increasingly rely on direct immune activation to limit pathogen burden, which could rapidly escalate the immunopathological risk. However, we stress that currently there is no evidence for this hypothesis in *T. molitor*.

Our work has important implications for the physiological mechanisms of ageing in insects. Our data reveal a baseline reduction in MT functioning with age, even in the absence of immune activation. This may indicate an age-dependent trade-off between investment in immune function and prevention of its immunopathological costs. It is possible that such increases in immunopathological damage to vital organs (even without infection) could be a key feature of senescence. However, we lack direct empirical evidence for this hypothesis. Further manipulative experiments are thus necessary to test whether experimental suppression of inflammatory pathways (e.g. RNAi of PO response in insects) results in limited reduction of MT activity in older uninfected individuals, in turn reducing their mortality. We also found that after infection, MTs of older beetles almost stopped functioning (~95% reduction). We thus propose that older beetles succumb to infection faster than younger beetles due to the inability of damaged MTs to effectively maintain physiological homeostasis.

Finally, our work also has important implications for the evolution of maladaptive immune pathology in natural populations. Natural selection might be almost blind to self-damage associated with immune response, because the costs are usually paid at a later stage in life (21). From a public health perspective, such relaxed selection can pose a serious problem to an ageing human population, potentially exacerbating the autoimmune disease crisis worldwide (1, 41, 42). We suggest that intimate connections between immune action and immunopathology may drive these hallmarks of ageing. We hope that our observations encourage further empirical work for a deeper understanding of life history tradeoffs and fitness impacts associated with immune function and its immunopathological outcome. These are necessary to understand how and why maladaptive immunopathological features of an immune system have evolved.

## ACKNOWLEDGEMENTS

We are grateful to Dino McMahon, Caroline Zanchi, Saurabh Mahajan and Vrinda Ravi Kumar for feedback on the manuscript. We thank Jayjit Das, Arun Prakash, Caroline Zanchi and Dipendra Nath Basu for their help during experiments and data analysis. We acknowledge funding and support from the Centre for International Collaboration, Free University of Berlin; DAAD and SERB-DST Young Investigator Grant supplements to IK; the National Centre for Biological Sciences, India; a DST Inspire Faculty fellowship to DA; and European Research Council (EVORESIN) grant supplement to JR.

## REFERENCES

1. Bach J-F (2002) The effect of infections on susceptibility to autoimmune and allergic diseases. N Engl J Med 347(12):911–920.

2. Dye C, Scheele S, Dolin P, Pathania V, Raviglione M (1999) Global burden of tuberculosis: Estimated incidence, prevalence, and mortality by country. JAMA 282(7):677–686.

3. Bekker L-G, Moreira A-L, Bergtold A, Freeman S, Ryfell B, Kaplan G (2000) Immunopathologic effects of tumor necrosis factor alpha in murine mycobacterial infection are dose dependent. Infect Immun 68(12):6954–6961.

4. Urabe K, Aroca P, Tsukamoto K, Mascagna D, Palumbo A, Prota G, Hearing V-J (1994) The inherent cytotoxicity of melanin precursors: A revision. Biochim Biophys Acta - Mol Cell Res 1221(3):272–278

5. Nappi A-J, Vass E, Frey F, Carton Y (1995) Superoxide anion generation in Drosophila during melanotic encapsulation of parasites. Eur J Cell Biol 68(4):450–456.

6. Zhao P, Lu Z, Strand M, Jiang H (2008) Antiviral, antiparasitic, and cytotoxic effects of 5,6-dihydroxyindole (DHI), a reactive compound generated by phenoloxidase during insect immune response. Insect Biochem Mol Biol 141(4):520–529.

7. Finch C-E, Crimmins E-M (2004) Inflammatory exposure and historical changes in human life-spans. Science 305(5691):1736–1739.

8. Pursall E-R, Rolff J (2011) Immune responses accelerate ageing: Proof-of-principle in an insect model. PLoS One 6(5): e19972.

9. Davies S-A, Overend G, Sebastian S, Cundall M, Cabrero P, Dow J-A, Terhzaz S (2012) Immune and stress response “cross-talk” in the Drosophila Malpighian tubule. J Insect Physiol 58(4):488–497.

10. Pacheco C, Alevi K, Ravazi A, M-T-V-A Oliveira (2014) Review: Malpighian tubule, an essential organ for insects. Entomol Ornithol Herpetol Curr Res 3(02):2–4.

11. Cerenius L, Lee BL, Söderhäll K (2008) The proPO-system: pros and cons for its role in invertebrate immunity. Trends Immunol 29(6):263–71.

12. Sadd B-M, Siva-Jothy M-T (2006) Self-harm caused by an insect's innate immunity. Proc Biol Sci B 273(1600):2571–4.

13. De Gregorio E, Han S-J, Lee W-J, Baek M-J, Osaki T, Kawabata S, Lee B-L, Iwanaga S, Lemaitre B, Brey P-T (2002) An immune-responsive Serpin regulates the melanization cascade in *Drosophila*. Dev Cell 3(4):581–592.

14. Ayres J-S, Schneider D-S (2008) A signaling protease required for melanization in Drosophila affects resistance and tolerance of infections. PLoS Biol 6(12):2764–2773.

15. Deveale B, Brummel T, Seroude L (2004) Immunity and aging □: the enemy within □? Aging Cell 3(4):195–208.

16. Shanley D-P, Aw D, Manley N-R, Palmer D-B (2009) An evolutionary perspective on the mechanisms of immunosenescence. Trends immunol 30(7): 374–381.

17. Licastro F, Candore G, Lio D, Porcellini E, Colonna-Romano G, Franceschi C, Caruso C (2005) Innate immunity and inflammation in ageing: a key for understanding age-related diseases. Immun Ageing 2:8.

18. Zerofsky M, Harel E, Silverman N, Tatar M (2005) Aging of the innate immune response in *Drosophila melanogaster*. Aging Cell:103–108.

19. Khan I, Prasad NG (2013) The aging of the immune response in *Drosophila melanogaster*. J Gerontol A Biol Sci Med Sci 68(2):129–35.

20. Khan I, Prakash A, Agashe D (2015) Immunosenescence and the ability to survive bacterial infection in the red flour beetle *Tribolium castaneum*. J Anim Ecol 85:291–301.

21. Hamilton WD (1966) The moulding of senescence by natural selection. J Theor Biol 12(1):12–45.

22. Mueller L, Rose M (1996) Evolutionary theory predicts late-life mortality plateaus. Evolution 93(26):15249–15253.

23. Haine E, Moret Y, Siva-jothy M, Rolff J (2008) Antimicrobial defence and persistent infection in insects. Science 322(21):1257–1259.

24. Dobson A-J, Johnston PR, Vilcinskas A, Rolff J (2012) Identification of immunological expressed sequence tags in the mealworm beetle Tenebrio molitor. J Insect Physiol 58(12):1556–1561.

25. Johnston P-R, Rolff J (2015) Host and symbiont jointly control gut microbiota during complete metamorphosis. PLoS Pathog 11(11): e1005246.

26. Schmittgen T-D, Livak KJ (2008) Analyzing real-time PCR data by the comparative CT method. Nat Protoc 3(6):1101–1108.

27. Maddrell S-H, Overton J-A (1990) Transport in insect Malpighian tubules. Methods Enzymol 192(1958):617–632.

28. Neufeld D-S, Leader J-P (1998) Cold inhibition of cell volume regulation during the freezing of insect Malpighian tubules. J Exp Biol 201:2195–2204.

29. Nicolson S (1992) Excretory function in *Tenebrio molitor*: Fast tubular secretion in a vapour-absorbing insect. J Insect Physiol 38(2):139–146.

30. Wiehart U-I-M, Nicolson S-W, Eigenheer R-A, Schooley D-A (2002) Antagonistic control of fluid secretion by the Malpighian tubules of *Tenebrio molitor*: Effects of diuretic and antidiuretic peptides and their second messengers. J Exp Biol 501:493–501.

31. Zanchi C, Troussard J, Martinaud G (2011) Differential expression and costs between maternally and paternally derived immune priming for offspring in an insect. J Anim Ecol 80:1174–1183.

32. Swindell W-R (2010) Accelerated failure time models provide a useful statistical framework for ageing research. Exp Gerontol 44(3):190–200.

33. Patel K, Kay R, Rowell L (2006) Comparing proportional hazards and accelerated failure time models: an application in influenza. Pharm Stat 5(3):213–224.

34. Wang C, Li Q, Redden D-T, Weindruch R, Allison D-B (2004) Statistical methods for testing effects on “maximum lifespan.” Mech Ageing Dev 125(9):629–632.

35. Brandt S-M, Dionne M-S, Khush R-S, Pham L-N, Vigdal T-J, Schneider D (2004) Secreted bacterial effectors and host-produced eiger/TNF drive death in a *Salmonella*-infected fruit fly. PLoS Biol 2(12): e418.

36. Stout-Delgado H-W, Du W, Shirali A-C, Booth C-J, Goldstein D-R (2009) Aging promotes neutrophil-induced mortality by augmenting IL-17 production during viral infection. Cell Host Microbe 6(5):446–456.

37. Corby-Harris V, Habel K-E, Ali F-G, Promislow D-E-L (2007) Alternative measures of response to *Pseudomonas aeruginosa* infection in *Drosophila melanogaste*r. J Evol Biol 20(2):526–33.

38. Ramsden S, Cheung Y-Y, Seroude L (2008) Functional analysis of the *Drosophila* immune response during aging. Aging Cell 7(2):225–236.

39. Vasto S, Candore G, Balistreri C-R, Caruso M, Colonna-Romano G, Grimaldi M-P, Listi F, Nuzzo D, Lio D, Caruso C (2007) Inflammatory networks in ageing, age-related diseases and longevity. Mech Ageing Dev 128(1):83–91.

40. Belloni V, Faivre B, Guerreiro R, Arnoux E, Bellenger J, Sorci G (2010) Suppressing an anti-inflammatory cytokine reveals a strong age-dependent survival cost in mice. PLoS One 5(9): e12940.

41. Zaccone P, Fehervari Z, Phillips J-M, Dunne D-W, Cooke A (2006) Parasitic worms and inflammatory diseases. Parasite Immunol 28(10):515–523.

42. Finch C-E (2010) Evolution of the human lifespan and diseases of aging: roles of infection, inflammation, and nutrition. Proc Natl Acad Sci USA 107 Suppl 1:1718–1724.

